# Effects of social defeat stress and fluoxetine treatment on neurogenesis and behaviour in mice that lack zinc transporter 3 (ZnT3) and vesicular zinc

**DOI:** 10.1101/776633

**Authors:** Brendan B. McAllister, Angela Pochakom, Selena Fu, Richard H. Dyck

## Abstract

Depression is a leading cause of disability worldwide, in part because the available treatments are inadequate and do not work for many people. The neurobiology of depression, and the mechanism of action of common antidepressant drugs such as selective serotonin reuptake inhibitors (SSRIs), is not well understood. One mechanism thought to underlie the effects of these drugs is upregulation of adult hippocampal neurogenesis. Evidence indicates that vesicular zinc is required for modulation of adult hippocampal neurogenesis, at least under some circumstances. Vesicular zinc refers to zinc that is stored in the synaptic vesicles of certain neurons, including in the hippocampus, and released in response to neuronal activity. It can be eliminated from the brain by deletion of zinc transporter 3 (ZnT3), as is the case in ZnT3 knockout mice. Here, we examined the effects of repeated social defeat stress and subsequent chronic treatment with the SSRI fluoxetine on behaviour and neurogenesis in ZnT3 knockout mice. We hypothesized that fluoxetine treatment would increase neurogenesis and reverse stress-induced behavioural symptoms in wild type, but not ZnT3 knockout, mice. As anticipated, stress induced persistent depression-like effects, including social avoidance and anxiety-like behaviour. Fluoxetine decreased social avoidance, though the effect was not specific to the stressed mice, but did not affect anxiety-like behaviour. Surprisingly, stress increased the survival of neurons born 1 day after the last episode of defeat stress. Fluoxetine treatment also increased cell survival, particularly in wild type mice, though it did not affect proliferation. Our results did not support our hypothesis that vesicular zinc is required for the behavioural benefits of fluoxetine treatment. As to whether vesicular zinc is required for the neurogenic effects of fluoxetine, our results were inconclusive, warranting further investigation into the role of vesicular zinc in adult hippocampal neurogenesis.

## 1. INTRODUCTION

Depression is a leading cause of disability worldwide and, as such, has been the subject of intensive scientific research. Yet much remains unknown about how depression develops, as well as how experiences such as chronic stress, a risk factor for depression (Kendler & Gardner, 2016; Kendler, Karkowski, & Prescott, 1999), contribute to this process. Also unknown is how pharmacological treatments for depression – the most common being selective serotonin reuptake inhibitor (SSRI) drugs – work on the brain to ameliorate the disorder, and why certain individuals respond to these drugs while others do not. One key to unravelling the etiology of depression is to understand the long-term structural changes that occur in the depressed brain, as well as how these changes are reversed, or compensated for, by antidepressant treatments.

A component of neuroplasticity that has relevance to both depression and the effects SSRIs is adult neurogenesis. This refers to the process of generating new neurons in the mature brain (Altman, 1962; Altman & Das, 1965). Predominantly, adult neurogenesis takes place at two sites: the subventricular zone and the hippocampal dentate gyrus. The latter has been linked to the etiology of depression. In the dentate gyrus – specifically in the subgranular zone (SGZ) of the granule cell layer – new cells are generated by asymmetrically-dividing radial glial-like neural progenitor cells (Encinas, Vaahtokari, & Enikolopov, 2006). The newly generated cells, called amplifying progenitor cells, divide symmetrically to produce mostly immature neurons and a smaller number of glia (Pham, Nacher, Hof, & McEwen, 2003; Tanapat, Hastings, Rydel, Galea, & Gould, 2001). For the most part, the neurons migrate into the granule cell layer (Cameron, Wooley, McEwen, & Gould 1993). Some undergo apoptosis within days or weeks of being born (Dayer, Ford, Cleaver, Yassaee, & Cameron 2003; Sairanen, Lucas, Ernfors, Castrén, & Castrén, 2005; Tanapat et al., 2001). Others survive and become integrated into existing neural circuits (Kee, Teixeira, Wang, & Frankland, 2007; van Praag et al., 2002), affecting hippocampal function and, ultimately, behaviour (Aimone, Wiles, & Gage, 2009; Clelland et al., 2009; Sahay et al., 2011; Snyder, Kee, & Wojtowicz, 2001).

The effects of stress, depression, and SSRIs on hippocampal neurogenesis have been well-described, at least in non-human animals. While it is difficult, if not impossible, to truly replicate human depression in rodents, procedures that induce stress and model depression-like behaviour have been used to study these effects. Stress, either acute or chronic, has a generally suppressive effect on neurogenesis (Czéh et al, 2001, 2002; Gould, McEwen, Tanapat, Galea, & Fuchs, 1997; Mitra, Sundlass, Parker, Schatzberg, & Lyons 2006; Pham et al., 2003; Surget et al., 2011; Tanapat et al., 2001). Chronic treatment with SSRIs, such as fluoxetine, has the opposite effect, boosting neurogenesis in rats and mice (Encinas et al., 2006; Malberg, Eisch, Nestler, & Duman, 2000; Sairanen et al., 2005). Importantly, this seems to be required for some, though not all, of the behavioural effects of SSRIs in stressed mice (David et al., 2009; Santarelli et al., 2003; Surget et al., 2008, 2011; Wang, David, Monckton, Battaglia, & Hen, 2008). How these findings apply to humans is not entirely clear, but there is evidence that treatment with antidepressant drugs, either tricyclics or SSRIs, increases hippocampal neurogenesis in depressed people (Boldrini et al., 2012; but see Lucassen, Stumpel, Wang, & Aronica, 2010), and that depression is associated with smaller hippocampal volume (Cole, Costafreda, McGuffin, & Fu, 2011; McKinnon, Yucel, Nazarov, & MacQueen, 2009; Sheline, Wang, Gado, Csernansky, & Vannier, 1996) and fewer dentate granule cells (Boldrini et al., 2013).

Another factor that appears to be involved in modulating hippocampal neurogenesis is vesicular zinc. This describes zinc that is stored within synaptic vesicles in the axon terminals of certain neurons (Brown & Dyck, 2004; Pérez-Clausell & Danscher, 1985), from where it can be released in an activity-dependent manner (Aniksztejn, Charton, & Ben-Ari, 1987; Assaf & Chung, 1984; Howell, Welch, & Frederickson, 1984). Once in the synaptic cleft, zinc can exert signaling effects by binding to a plethora of targets, including many neurotransmitter receptors (McAllister & Dyck, 2017). Sequestration of zinc into synaptic vesicles is the function of a membrane transport protein called zinc transporter 3 (ZnT3) (Palmiter, Cole, Quaife, & Findley, 1996; Wenzel, Cole, Born, Schwartzkroin, & Palmiter, 1997). If this protein is eliminated, as is the case in ZnT3 knockout (KO) mice, vesicular zinc is undetectable in the brain (Cole, Wenzel, Kafer, Schwartzkroin, & Palmiter, 1999). Evidence suggests that, under certain conditions, ZnT3 KO mice exhibit abnormalities in adult hippocampal neurogenesis. Whereas neurogenesis is upregulated in response to hypoglycemia in wild type (WT) mice, the same effect is not observed in ZnT3 KO mice (Suh et al., 2009). Unpublished experiments from our laboratory provide further evidence. Normally, neurogenesis can be enhanced in rodents by transferring them from standard laboratory housing to more complex, stimulating environments (Kempermann, Brandon, & Gage, 1997; Kempermann, Kuhn, & Gage, 1998, or, as mentioned above, by treating them with SSRIs. However, in our experiments, ZnT3 KO mice did not show enhanced neurogenesis in response to enriched housing (Chrusch, 2015) or treatment with fluoxetine (Boon, 2016). Furthermore, they did not exhibit the behavioural benefits (i.e., enhanced spatial cognition) of enriched housing (Chrusch, 2015).

Previously, we studied the effects of subjecting ZnT3 KO mice to repeated social defeat (RSD) stress (McAllister, Wright, Wortman, Shultz, & Dyck, 2018), which results in behavioural changes that reflect some aspects of depression (Krishnan et al., 2007). These mice exhibited some of the same effects of stress as WT mice, including increased anxiety-like behaviour and social avoidance of an aggressive CD-1 mouse. However, unlike WT mice, ZnT3 KO mice did not become socially avoidant of a same-strain conspecific, suggesting that ZnT3 KO mice are less susceptible to social avoidance following RSD stress. The present study sought to extend these results, with two main objectives. Given that vesicular zinc may be required for modulation of neurogenesis in response to certain experiences, we first sought to assess if it is also required for modulation of neurogenesis in response to RSD. If so, this could help to explain our finding that ZnT3 KO mice are less susceptible to stress. Second, given that neurogenesis has been implicated in the antidepressant effects of SSRIs, we sought to test whether stressed ZnT3 KO mice would show behavioural benefits from chronic treatment with the SSRI fluoxetine. Our hypothesis, based on our previous findings, was that fluoxetine would increase neurogenesis in WT mice, and that this would be associated with reduced social avoidance of a CD-1 mouse and reduced anxiety-like behaviour, whereas ZnT3 KO mice would not exhibit increased neurogenesis, and would show no behavioural benefits from fluoxetine treatment.

## 2. METHOD

### 2.1. Animals

All protocols were approved by the Life and Environmental Sciences Animal Care Committee at the University of Calgary and followed the guidelines for the ethical use of animals provided by the Canadian Council on Animal Care. Mice were housed in temperature- and humidity-controlled rooms maintained on a 12:12 light/dark cycle (lights on during the day). Food and water were provided *ad libitum* except where otherwise noted. WT and ZnT3 KO mice, on a mixed C57BL/6×129/Sv genetic background, were bred from heterozygous pairs. Offspring were housed with both parents until P21, at which point they were weaned and housed in standard cages (28 × 17.5 × 12 cm with bedding, nesting material, and one enrichment object) in groups of 2-5 same-sex littermates. CD-1 mice used for the RSD procedure were retired breeders, 4-12 months old, acquired from the University of Calgary Transgenic Services Facility or from Charles River. CD-1 mice were single-housed in standard cages except as described below.

### 2.2. Experimental Design and BrdU Administration

For a diagram depicting the experimental design, see Figure 1. At 8-10 weeks of age, WT and ZnT3 KO mice were assigned to one of four treatment conditions: control + vehicle (WT: *n* = 11; KO: *n* = 9); control + fluoxetine (WT: *n* = 10; KO: *n* = 10); stress + vehicle (WT: *n* = 10; KO: *n* = 9); stress + fluoxetine (WT: *n* = 10; KO: *n* = 10). The stress consisted of 10 days of RSD (day 1 to 10), followed by isolated housing for the remainder of the experiment. The control mice remained in standard housing (described in section 2.1) throughout the experiment.

**Figure 1.**
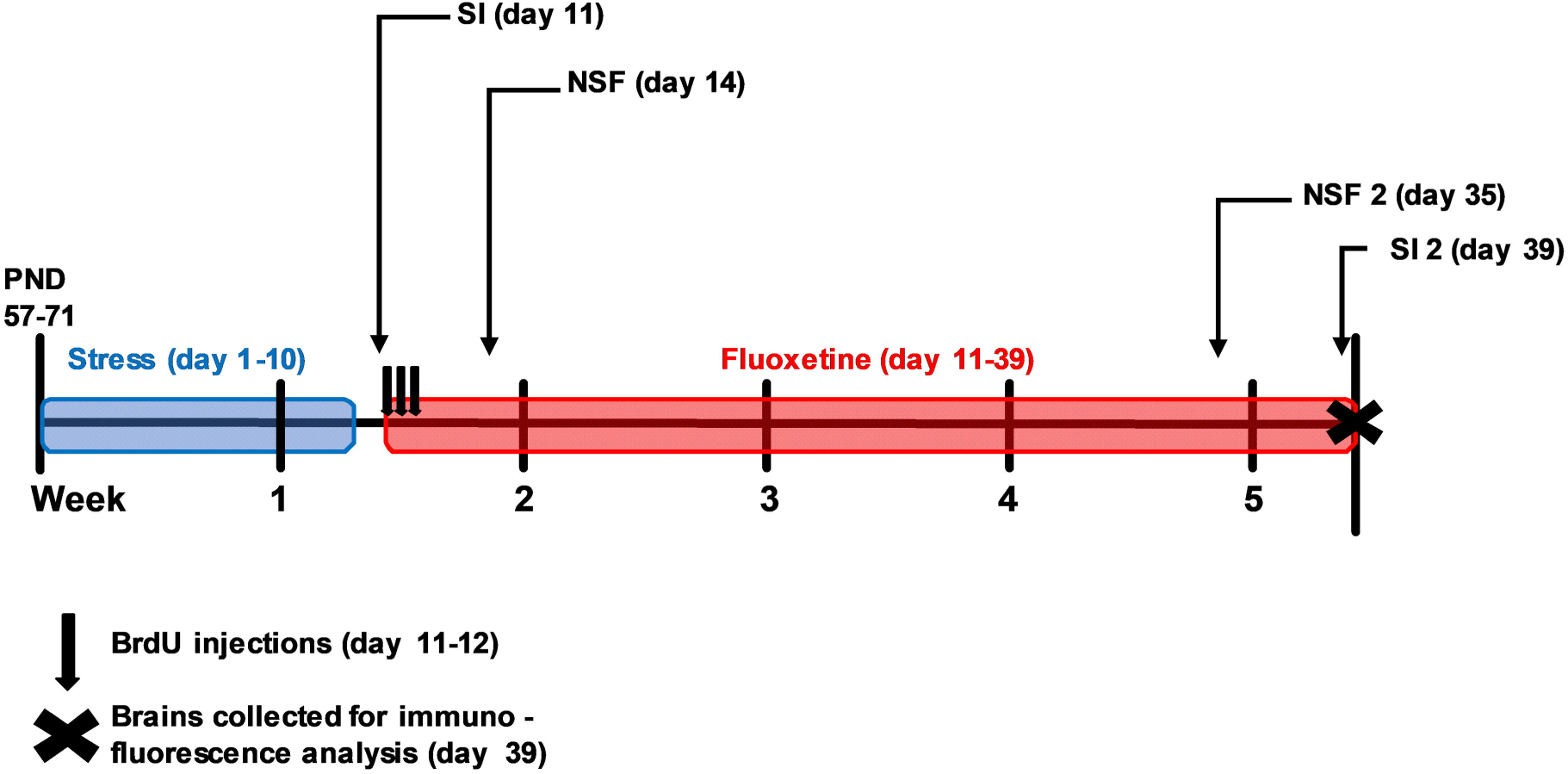
Timeline depicting the experimental design. WT and ZnT3 KO mice, 8-10 weeks old (postnatal day 57-71) at the beginning of the experiment, were subjected to 10 days of repeated social defeat (RSD) stress, consisting of daily episodes of social defeat, followed by isolated housing for the remainder of the experiment. Control WT and ZnT3 KO mice remained in group housing with their same-sex littermates throughout the experiment. Beginning 1-day post-RSD, and lasting for 28 days, some mice received fluoxetine (25 mg/kg/day) in their drinking water. This procedure resulted in eight treatment groups (*n* = 9-11).

One-day post-stress (day 11), mice were subjected to the first social interaction test. Immediately following this test, the first injection of 5-bromo-2 •-deoxyuridine (BrdU; Sigma) was administered to label cells in S-phase of the cell cycle. In total, three injections (100 mg/kg, IP) of BrdU dissolved in sterile-filtered PBS (15 mg/ml) were administered at 8-h intervals. Also immediately following the social interaction test, the mice began 4 weeks of fluoxetine treatment. Four-days post-stress (day 14), the mice were subjected to the novelty-suppressed feeding (NSF) test. The NSF test was re-administered 3 weeks later (day 35), after 24 d of fluoxetine treatment. Finally, the social interaction test was re-administered after the 4 weeks of fluoxetine treatment was completed (day 39). Following this, the mice were killed and brains were extracted and processed for immunofluorescence labeling.

### 2.3 Repeated Social Defeat

The RSD procedure was adapted from Golden, Covington, Berton, and Russo (2011). Mice were subjected to daily episodes of defeat over 10 days. For each defeat, the mouse was transferred to a novel CD-1 mouse’s cage for a period of 5 min. During this time, the CD-1 resident would reliably attack the smaller intruder. After three attacks (each defined as an uninterrupted episode of physical interaction, almost always resulting in vocalizations from the intruder) the intruder was placed in a cylindrical mesh enclosure (8.5 cm?) for the remainder of the 5 min, allowing the mice to interact in close proximity but restricting further fighting or injury. Following this, the intruder was housed with the CD-1 resident that had just defeated it, with the two mice separated by a perforated acrylic partition dividing the large cage (24 × 45.5 × 15 cm) lengthwise into two compartments, allowing for visual, auditory, and olfactory contact between the mice, but limiting physical interaction. Prior to each defeat session, the intruder mice were rotated between cages, in order to prevent them from habituating to a particular CD-1 resident. After the final defeat, the mice were returned to single-housing in standard cages for the remainder of the experiment.

To ensure that the CD-1 residents would reliably engage with and defeat the intruders, CD-1 mice were screened for aggressiveness prior to the experiments. The screening procedure consisted of three trials (one per day for 3 days) in which an intruder was introduced into the cage of the CD-1 resident for 3 min. (The intruders used for screening were ZnT3 heterozygotes, retired from previous experiments or surplus from our breeding colony.) Only CD-1 mice that attacked the intruder within 30 s on two or more consecutive trials were used for the experiment. To further promote territoriality and aggression, the CD-1 mice were housed in large cages for at least 24 h prior to the introduction of the first intruder, and remained in the same cages throughout the 10-day procedure.

### 2.4. Fluoxetine Treatment

Fluoxetine hydrochloride (Santa Cruz) was administered orally (approximately 25mg/kg/day) via the drinking water, as previously described (McAllister, Kiryanova, & Dyck, 2012). Fluoxetine was dissolved in distilled water (1 mg/ml) and then diluted to the appropriate concentration in tap water. To achieve the correct dosage, mice were weighed, and the amount of water consumed over the previous 3-4 days was measured using calibrated water bottles. In cages with multiple mice (i.e., group-housed controls), the average weight of the mice was used to calculate dosages. The dosage was adjusted, and fresh fluoxetine solution provided, two times per week. Vehicle-treated mice were weighed and handled on the same schedule but received only tap water.

### 2.5. Behavioural Assessment

All testing was conducted during the light phase of the mice’s light/dark cycle, and mice were habituated to the testing room for at least 30 min prior to the start of each test.

#### 2.5.1 Social interaction

The procedure for the social interaction test was adapted from Golden et al. (2011). The test was conducted under dim red light. The apparatus for the test was an open field (40 × 40 cm) constructed of white corrugated plastic. The test consisted of three 150 s phases, each separated by 60 s. For the first phase, a cylindrical mesh enclosure (10 cm?, with a weighted glass flask fixed to the roof to prevent mice from climbing on top of the enclosure or moving the enclosure) was placed against a wall of the field; the mouse was then placed along the center of the opposing wall and allowed to explore freely. The second phase was the same, but with a novel, age-matched mouse of the same strain (novel conspecific) placed inside the enclosure. For the third phase, the conspecific was replaced by a novel, aggressive CD-1 mouse. Between trials, the mouse being tested was returned to its home cage. A different but identical mesh enclosure was used for the first phase versus the second and third phases, in order to reduce odours on the enclosure that might influence exploratory behaviour. Between testing each mouse, the enclosures and the field were cleaned with Virkon; the field was also cleaned of urine and feces between each phase. The test was recorded using a digital video camera with night-vision capability (Sony HDR-SR8), and scoring was automated using tracking software (ANY-maze, version 4.73). The following parameters were scored: “interaction time” (i.e., time spent in the interaction zone, defined as a 26 × 16 cm rectangle around the mesh enclosure); “corner time” (i.e., time spent in either of the two corners of the field opposing the enclosure, with each corner encompassing a 9 × 9 cm area); and total distance traveled. Social interaction ratios were calculated by dividing interaction time with the CD-1 mouse during the third phase by interaction time with the empty enclosure during the first phase. Based on the interaction ratio in the first social interaction test, mice were defined as susceptible (interaction ratio < .5) or resilient (interaction ratio • 0.5), following the definitions used in our previous work (McAllister et al., 2018).

#### 2.5.2 Novelty-suppressed feeding

In the NSF test, the mouse faces a conflict between two motivations: the drive to feed when hungry and the inclination to avoid feeding in an exposed environment. The latency to feed in this test provides an indicator of anxiety-like behaviour, with longer latencies assumed to reflect greater anxiety. The protocol was adapted from Samuels and Hen (2011). Mice were food deprived for 16 h prior to the test. The test was conducted in an open field under bright lighting (∼800 lux). The floor of the field was covered with wood-chip bedding. (The field was not entirely novel to the mice, as it was the same apparatus used for the social interaction test, but it did include some novel features, such as the bedding on the floor, the room in which the field was situated, and the lighting of the room.) A food pellet (standard mouse chow) was fixed to a small platform in the center of the field with an elastic band, preventing the mouse from moving the pellet. The latency to begin feeding was recorded, up to a maximum time of 10 min, at which point the test was terminated and the mouse was assigned a latency of 600 s. The mouse was then returned to its home-cage (with its cage-mates temporarily removed), and transported immediately to an adjacent, dimly lit (∼3 lux) room. A pre-weighed food pellet was placed in the food hopper, and the latency to begin feeding in the home cage was recorded; a maximum latency score of 180 s was given if the latency to feed exceeded that length. After the mouse began feeding, it was allowed 5 min to eat, after which the pellet was removed and weighed to calculate food consumption. Body weight was also recorded both prior to food deprivation and after the test.

### 2.6. Tissue Preparation and Immunofluorescence Labeling

Mice were deeply anaesthetized with an overdose of sodium pentobarbital, and transcardially perfused with phosphate buffered saline (PBS) until the blood was cleared, followed by perfusion with 4% paraformaldehyde (PFA) in PBS. Brains were extracted and post-fixed overnight in 4% PFA in PBS at 4 °C. After post-fixing, the brains were transferred to a sucrose solution (30% sucrose, 0.02% sodium azide in PBS) and stored at 4°C. Brains were cut coronally into six series of 40 µm sections using a freezing, sliding microtome (American Optical, Model #860). Immunofluorescence labeling was conducted for markers of neurogenesis. One series of tissue sections was labeled for the cellular proliferation marker Ki67 (primary antibody: anti-Ki67, Abcam, Cat# ab15580, RRID: AB_443209, 1:2000, 0.5 • g/ml). A second series was labeled for BrdU to assess cell survival (primary antibody: anti-BrdU, Bio-Rad, Cat# MCA2060, RRID: AB_323427, 1:200). Following immunolabeling, sections were mounted on gelatin-coated slides, coverslipped with fluorescence mounting medium, and stored at 4 °C until analysis. Detailed immunofluorescence protocols are provided in the Supplemental Methods.

### 2.7. Cell Counting

Ki67^+^ and BrdU^+^ cells were counted using an epi-fluorescence microscope (Zeiss Axioskop 2) with a 63×/1.40 objective. Cells were counted in the granule cell layer and the SGZ (defined as three cell-widths from the hilar edge of the granule cell layer) in all sections containing the hippocampal dentate gyrus. The counts were then multiplied by six (since the brains were sectioned into six series) to estimate the total number of cells.

### 2.8. Statistical Analysis

Statistical analyses were conducted using IBM SPSS Statistics (version 24). Unless otherwise stated, comparisons were conducted by three-way analysis of variance (ANOVA) with genotype (WT vs. ZnT3 KO), stress (control vs. stress), and drug (vehicle vs. fluoxetine) as factors. Significant interactions were followed-up with simple effects tests using the pooled error term, unless equality of variances could not be assumed (Levene’s test: *p* < .05), in which case non-pooled error terms were used. All ANOVA results are reported in Supplemental Tables S1 and S2.

## 3. RESULTS

### 3.1. Neurogenesis

#### 3.1.1. Cell survival

One-day post-RSD, we injected mice with BrdU to label newborn cells. Brains were collected 4 weeks later, and the number of surviving cells was assessed by counting BrdU^+^ cells in the granule cell layer and SGZ of the hippocampal dentate gyrus (Figure 2A). Stress had a significant effect on the number of BrdU^+^ cells [*F*(1,71) = 21.88, *p* < .001]. Contrary to our hypothesis, however, the effect was not a decrease but rather an increase of 37% (Figure 2B). Further contradicting our hypothesis, the effect of stress on the number of BrdU^+^ cells was not limited to the WT mice [stress × genotype interaction: *F*(1,71) = 0.84, *p* = .362]. In summary, stress increased the number of cells that were born 1 day post-stress and survived at least 4 weeks, and this modulation of hippocampal neurogenesis was not dependent on vesicular zinc.

**Figure 2.**
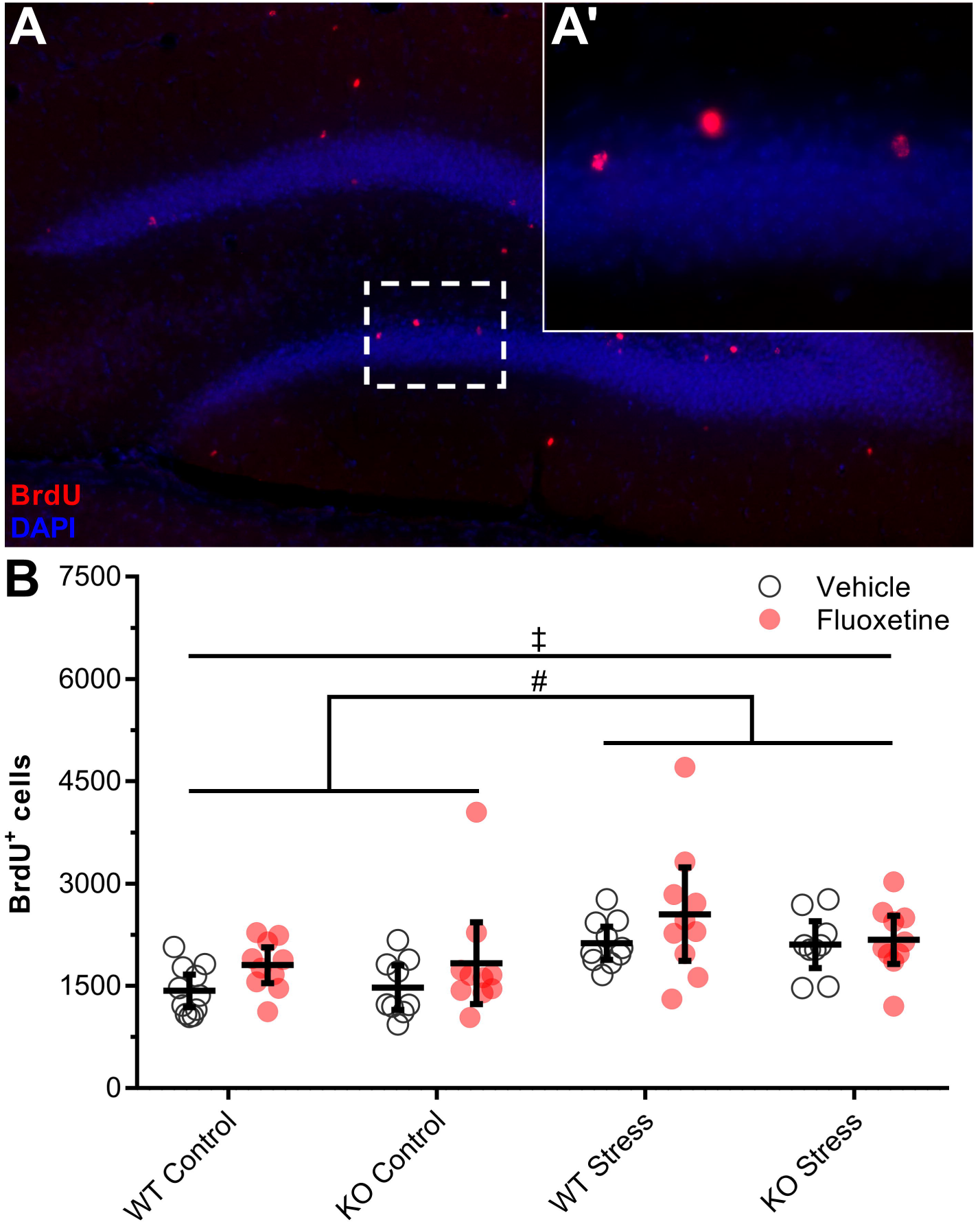
Hippocampal neurogenesis in WT and ZnT3 KO mice following repeated social defeat (RSD) stress and chronic fluoxetine treatment. BrdU was administered 1-day after the final episode of RSD stress to label dividing cells, and brains were collected 4 weeks later to examine their survival. BrdU^+^ cells were counted in the granule cell layer and subgranular zone of the dentate gyrus. (**A**) Sample image of the hippocampal dentate gyrus immunolabeled for BrdU. (**A•**) High magnification image of BrdU^+^ cells. (**B**) Stress significantly increased the number of BrdU^+^ cells that survived up to 4 weeks. Fluoxetine treatment during the 4-week period also increased the number of BrdU^+^ cells. As hypothesized, fluoxetine had a significant effect in WT mice but not in ZnT3 KO mice, though there was not a significant genotype × fluoxetine interaction. Error bars represent 95% confidence intervals (CIs). ^#^effect of stress, *p* < .05; ^‡^effect of fluoxetine, *p* < .05

We used planned contrasts to test our *a priori* hypothesis that 4 weeks of fluoxetine treatment would promote neurogenesis in WT mice but have no effect on ZnT3 KO mice. Consistent with our hypothesis, we found that fluoxetine increased the number of BrdU^+^ cells in WT mice [*F*(1,71) = 5.00, *p* = .028] while having no significant effect on ZnT3 KO mice [*F*(1,71) = 1.31, *p* = .256]. However, an ANOVA did not show that the effect of fluoxetine treatment differed significantly between genotypes [genotype × drug interaction: *F*(1,71) = 0.53, *p* = .470]. Instead, there was a significant main effect of fluoxetine treatment [*F*(1,71) = 5.64, *p* = .020], with fluoxetine increasing the number of BrdU^+^ cells by 18%. Thus, our results unambiguously support an effect of fluoxetine treatment on neurogenesis in WT mice, but did not provide clear support for our hypothesis that the neurogenic effect of fluoxetine would be dependent on vesicular zinc

We also examined whether the effect of stress on the number of BrdU^+^ cells differed between mice that were behaviourally susceptible or resilient to stress, based on the first social interaction test. A one-way ANOVA [*F*(2,73) = 12.14, *p* < .001] with Fisher’s LSD post-hoc test showed that, relative to controls, the number of BrdU^+^ cells was increased both in susceptible (*p* < .001) and resilient mice (*p* = .001). Susceptible and resilient mice did not differ (*p* = .871) (data not shown).

#### 3.1.2. Proliferation

In addition to the number of surviving cells, the number of proliferating (Ki67^+^) cells in the granule cell layer and SGZ of the hippocampal dentate gyrus was assessed (Figure 3A). First, we examined whether RSD stress, experienced 4 weeks before brains were collected, had an effect on proliferation. We found that it did not [main effect of stress: *F*(1,71) = 0.09, *p* = .769; Figure 3B]. There was also no effect of genotype on proliferation [*F*(1,71) = 0.11, *p* = .747]. Next, we used planned contrasts to test our *a priori* hypothesis that fluoxetine would increase proliferation in WT mice but not in ZnT3 KO mice.

**Figure 3.**
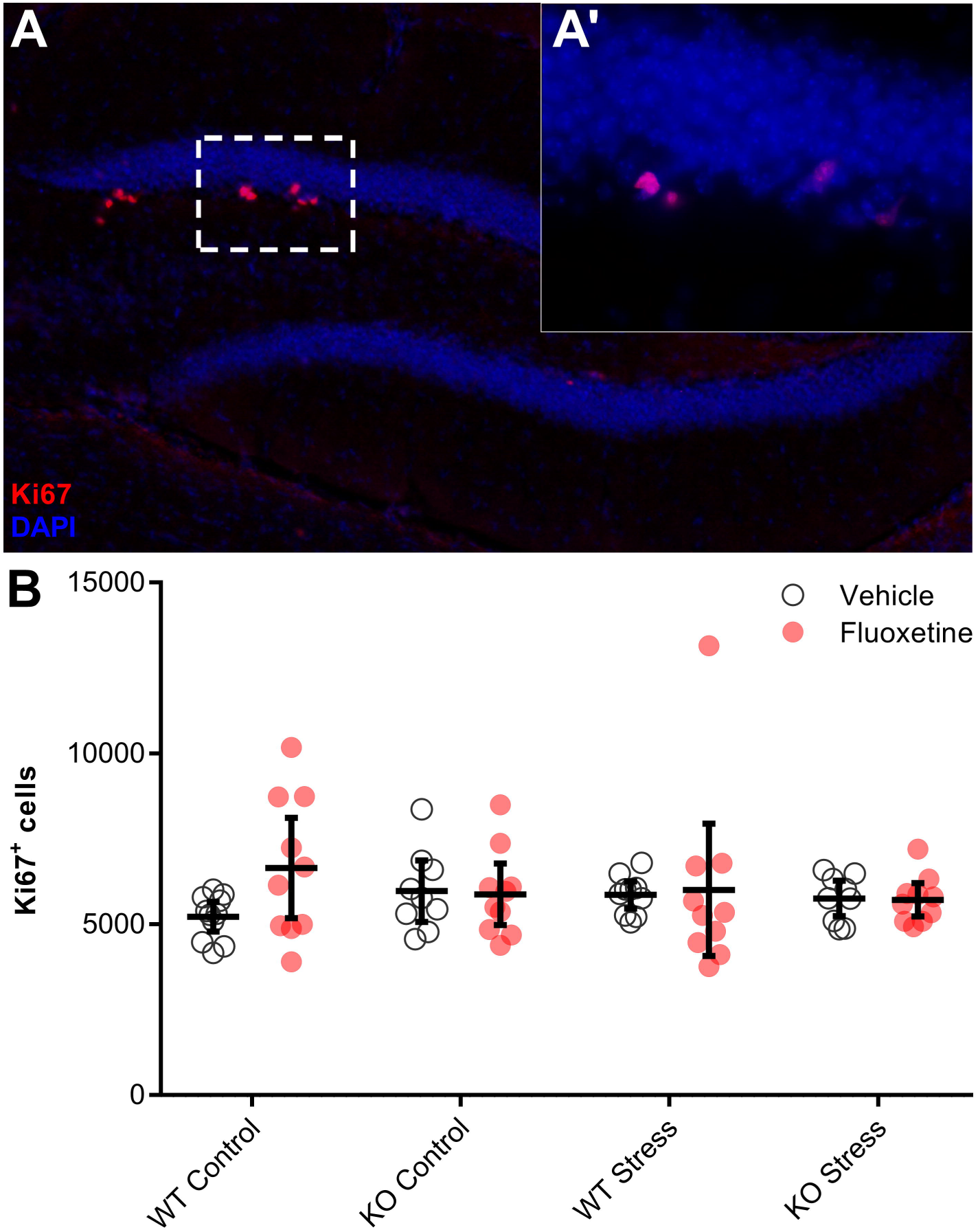
Hippocampal cell proliferation in WT and ZnT3 KO mice following repeated social defeat (RSD) stress and chronic fluoxetine treatment. Mice were subjected to 10 days of RSD stress, followed by 4 weeks of fluoxetine treatment, after which brains were collected and cell proliferation was assessed by counting the number of Ki67^+^ cells in the granule cell layer and subgranular zone of the dentate gyrus. (**A**) Sample image of the hippocampal dentate gyrus immunolabeled for Ki67. (**A•**) High magnification image of Ki67^+^ cells. (**B**) There was no effect of stress, fluoxetine treatment, or genotype on the number of proliferating cells. Error bars represent 95% CIs.

Fluoxetine had no significant effect on proliferation in either the WT mice [*F*(1,39) = 2.24, *p* = .157; Welch’s test] or in the ZnT3 KO mice [*F*(1,36) = 0.04, *p* = .834]. Furthermore, an ANOVA confirmed that the effect of fluoxetine treatment did not differ significantly between genotypes [genotype × drug interaction: *F*(1,71) = 1.77, *p* = .188]; nor was there a main effect of fluoxetine treatment [*F*(1,71) = 1.26, *p* = .265]. Finally, we assessed the effect of fluoxetine independently for each of the conditions. Fluoxetine treatment had no significant effects on proliferation [KO-control: *F*(1,18) = 0.03, *p* = .867; WT-stress: *F*(1,17) = 0.03, *p* = .871; KO-stress: *F*(1,17) = 0.02, *p* = .900], though it did tend to increase proliferation in the WT-control mice [*F*(1,19) = 4.80, *p* = .041; Welch’s test]. In summary, the results did not provide a good test of our hypothesis that the neurogenic effect of fluoxetine treatment is dependent on vesicular zinc, due to the lack of a robust effect of fluoxetine treatment on cell proliferation even in the WT mice.

### 3.2. Behaviour

#### 3.2.1. Social interaction

##### 3.2.1.1. First interaction test: replication of previous results

Previously, we found that RSD stress caused WT mice to become socially avoidant of both novel conspecifics and aggressive CD-1 mice, whereas stressed ZnT3 KO mice avoided only the CD-1 aggressors (McAllister et al., 2018). Here, we examined whether we could replicate this effect in a different sample of mice. We examined behaviour in the social interaction test, 1 day after the final episode of RSD. Because this test was conducted before the start of fluoxetine treatment, we were able to combine the vehicle- and fluoxetine-treated groups for analysis. We conducted planned contrasts to test the *a priori* hypotheses, based on our previous results, that in the test with a novel conspecific 1) interaction time and corner time would not differ between control and stressed ZnT3 KO mice, and 2) interaction time would be significantly decreased, and corner time significantly increased, in stressed WT mice relative to controls. We further hypothesized that in the social interaction test with an aggressive CD-1 mouse there would be significant main effects of stress on interaction time and corner time, with stress decreasing interaction time and decreasing corner time.

We first examined interaction with a novel conspecific (Figure 4A). Consistent with our hypotheses, stress decreased interaction time in the WT mice by 32% [*F*(1,39) = 5.15, *p* = .024], while having no significant effect on the ZnT3 KO mice [*F*(1,36) = 0.70, *p* = .410; Welch’s test for unequal variances]. However, a two-way ANOVA showed no significant interaction between the effects of genotype and stress [*F*(1,75) = 1.36, *p* = .248]. Time spent in the corners of the field exhibited a similar pattern. Planned contrasts showed that stress had a significant effect on the WT mice, more than doubling corner time [*F*(1,39) = 5.26, *p* = .034; Welch’s test], while having no significant effect on the ZnT3 KO mice [*F*(1,36) = 3.72, *p* = .062; Welch’s test], though stress did tend to increase corner time in these mice. Again, a two-way ANOVA revealed no significant interaction between the effects of genotype and stress [*F*(1,75) = 0.33, *p* = .568]. Due to the lack of significant interaction effects, the results did not fully replicate our previous finding that the effect of RSD stress on social interaction with a novel conspecific is less pronounced in ZnT3 KO mice compared to WT mice.

**Figure 4.**
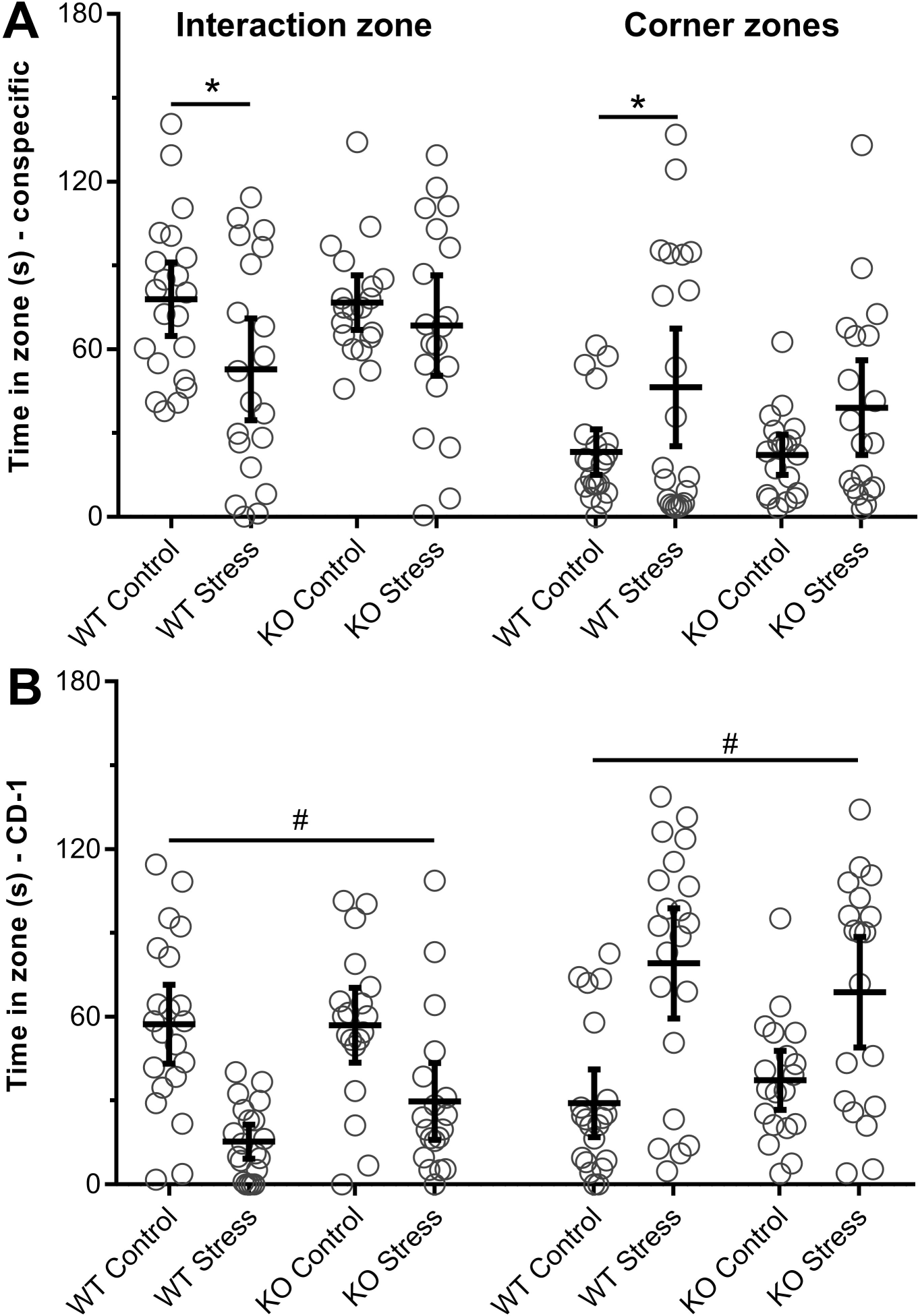
Results of the first social interaction test, conducted 1-day after the final episode of repeated social defeat stress. Because this test was conducted prior to the onset of fluoxetine treatment, groups were collapsed across this variable. (**A**) Interaction with a novel same-strain conspecific. As hypothesized, stress decreased interaction time and increased corner time in WT mice but had no significant effect on ZnT3 KO mice. However, there was not a significant genotype × stress interaction for either interaction time or corner time. (**B**) Interaction with a novel, aggressive CD-1 mouse. Stress decreased time spent in the interaction zone and increased time spent in the corner zones. Error bars represent 95% CIs. ^#^main effect of stress, *p* < .05; *difference between groups, *p* < .05

Next, we examined behaviour during the social interaction test with a novel, aggressive CD-1 mouse (Figure 4B). For interaction time, a two-way ANOVA confirmed our hypothesis that there would be a significant main effect of stress [*F*(1,75) = 34.70, *p* < .001] but no effect of genotype [*F*(1,75) = 1.42, *p* = .238] or interaction between the two factors [*F*(1,75) = 1.59, *p* = .212]. Stress decreased interaction time by 61%. For corner time, a two-way ANOVA also confirmed our hypothesis that there would be a significant main effect of stress [*F*(1,75) = 30.18, *p* < .001], but no effect of genotype [*F*(1,75) = 0.10, *p* = .750], or interaction [*F*(1,75) = 1.93, *p* = .169]. In the presence of a CD-1 mouse, stress more than doubled corner time. Finally, stress decreased the interaction ratio with a CD-1 mouse [*F*(1,75) = 14.71, *p* < .001], and there was no significant difference between genotypes [*F*(1,75) = 3.23, *p* = .074], though WT mice tended to have lower interaction ratios than ZnT3 KO mice (Table 1). Together, these results indicate that, as predicted, RSD stress caused avoidance of a novel, aggressive CD-1 mouse in both WT and ZnT3 KO mice.

**Table 1.** Additional behavioural measures from the first social interaction test. The test was conducted prior to the start of fluoxetine treatment. Statistics are reported as mean ± standard deviation. ^#^Main effect of stress, *p* < .05

|  | Wild type control | Wild type stress | ZnT3 KO control | ZnT3 KO stress |
| --- | --- | --- | --- | --- |
| **Social interaction test** | ($n = 21$) | ($n = 20$) | ($n = 19$) | ($n = 19$) |
| Distance (m) – empty cage | $8.7 \pm 1.8$ | $8.1 \pm 2.1$ | $9.1 \pm 2.0$ | $8.6 \pm 1.4$ |
| Interaction time (s) – empty cage<sup>#</sup> | $72.5 \pm 18.4$ | $62.0 \pm 24.2$ | $67.2 \pm 19.0$ | $51.1 \pm 20.0$ |
| Interaction ratio – CD-1<sup>#</sup> | $0.84 \pm 0.50$ | $0.31 \pm 0.31$ | $0.97 \pm 0.63$ | $0.60 \pm 0.60$ |

In the first phase of the social interaction test (empty holding cage), stress had no effect on total distance traveled [*F*(1,75) = 1.56, *p* = .216], nor was there a difference between genotypes (Table 1). There was, however, a significant effect of stress on time spent in the interaction zone, consistent with our previous results (McAllister et al., 2018). Stress decreased interaction zone time by 19% [*F*(1,75) = 8.28, *p* = .005]. There was no significant difference between genotypes [*F*(1,75) = 3.07, *p* = .084], though interaction time with the empty holding cage tended to be lower in ZnT3 KO mice than in WT mice (Table 1).

##### 3.2.1.2 Second interaction test: effect of fluoxetine treatment

Four weeks after the first test, we examined social interaction behaviour a second time to assess how stress-induced social avoidance was affected by chronic fluoxetine treatment. We focused here on interaction with an aggressive CD-1 mouse, as this was clearly decreased in both genotypes on the first interaction test, providing a similar baseline from which to assess the effects of fluoxetine treatment. Our hypotheses were that 1) in stressed mice that received vehicle treatment, social avoidance would persist to the second test (i.e., the mice would exhibit decreased interaction time and increased corner time relative to controls); and 2) fluoxetine treatment would reduce social avoidance (i.e., increase interaction time and decrease corner time) in stressed WT mice, but not in stressed ZnT3 KO mice.

We first examined time spent in the interaction zone (Figure 5A). In the second social interaction test, there was no longer a significant main effect of stress on interaction time [*F*(1,71) = 2.80, *p* = .098]. This did not appear to be due to a reversal in social avoidance specific to the fluoxetine-treated mice, however [stress × drug interaction: *F*(1,71) = 0.13, *p* = .716]; both the stressed fluoxetine-treated and stressed vehicle-treated mice showed similar amounts of interaction time, and there was no significant main effect of fluoxetine treatment [*F*(1,71) = 0.71, *p* = .403]. Furthermore, a simple-effects test showed that there was no effect of stress even when the comparison was limited to the vehicle-treated mice [*F*(1,71) = 0.84, *p* = .362]. This suggests that, even without drug treatment, social avoidance diminished over the 4 weeks between the first and second test. Our first hypothesis was thus not confirmed. Because there was no persistent effect of stress in the vehicle-treated mice, these results could not provide an adequate test of our second hypothesis.

**Figure 5.**
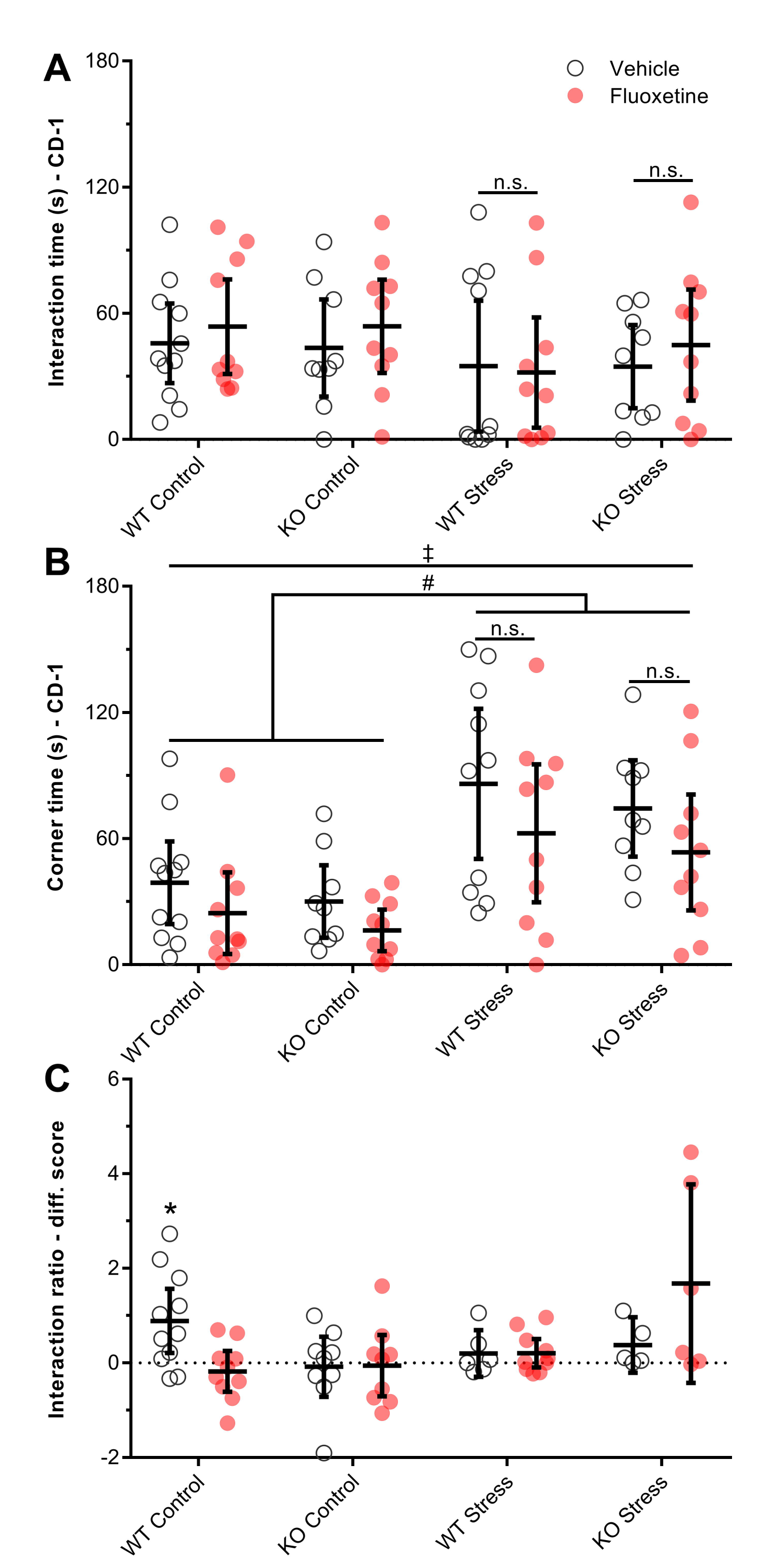
Results of the second social interaction test, conducted after 4 weeks of fluoxetine treatment. (**A**) Interaction time with a novel, aggressive CD-1 mouse. Contrary to our hypothesis, there was no persistent effect of stress on interaction time. Fluoxetine treatment also had no effect on stressed WT or ZnT3 KO mice. (**B**) Corner time during the interaction test with a novel, aggressive CD-1 mouse. The stressed mice exhibited more corner time, indicating that the effect of stress on social avoidance persisted at least 4 weeks. Fluoxetine treatment decreased corner time, indicating decreased social avoidance, though this effect was not limited to the stressed mice. (**C**) Comparison of social interaction behaviour in the first and second tests. Difference scores were calculated; a positive score indicates an increase in social interaction over time. Social interaction was defined using the interaction ratio, calculated by dividing the time spent interacting with a novel, aggressive CD-1 mouse (phase 3 of the test) by the time spent interacting with an empty holding cage (phase 1 of the test). With one exception (WT-control-vehicle group), interaction ratios were stable over time at the group level, as indicated by scores that did not differ significantly from zero. There was no effect of stress, fluoxetine treatment, or genotype on this measure. Error bars represent 95% CIs. ^#^effect of stress, *p* < .05; ^‡^effect of fluoxetine, *p* < .05; *difference from zero; n.s. indicates *p* > .05

We next examined time spent in the corners of the field (Figure 5B). Unlike interaction time, there was a significant main effect of stress on corner time [*F*(1,71) = 29.27, *p* < .001], with the stressed mice spending more than twice as much time in the corners as the control mice. Simple effects tests showed that this was also the case when the comparison was limited to vehicle-treated mice [*F*(1,37) = 17.34, *p* < .001; Welch’s test] and when it was further broken down by genotype [WT: *F*(1,19) = 7.12, *p* = .021, Welch’s test; KO: *F*(1,16) = 12.71, *p* = .003], supporting our first hypothesis. Regarding our second hypothesis, planned contrasts showed that there was no effect of fluoxetine treatment on the stressed WT mice [*F*(1,35) = 1.57, *p* = .218] or the stressed ZnT3 KO mice [*F*(1,35) = 1.17, *p* = .286]. Thus, our hypothesis that fluoxetine treatment would decrease corner time in WT mice but not in ZnT3 KO mice was not confirmed. There was, however, a significant main effect of fluoxetine treatment [*F*(1,71) = 5.59, *p* = .021], with fluoxetine decreasing the amount of time spent in the corners by 31%. This was not specific to the stressed mice [stress × drug: *F*(1,71) = 0.28, *p* = .600], and simple-effects tests showed that, when control and stressed mice were compared separately, there was no effect of fluoxetine treatment on either the control [*F*(1,36) = 3.42, *p* = .073] or stressed mice [*F*(1,35) = 2.73, *p* = .108]. To summarize, stressed mice still exhibited increased corner time 4 weeks after the final episode of RSD, and fluoxetine treatment reduced corner time globally, rather than having an antidepressant-like effect that was specific to the stressed mice.

One possible explanation for the lack of effect of fluoxetine on the stressed mice was that we included in our analysis both mice that were susceptible to stress and mice that were resilient. Typically, susceptibility in the social interaction test is defined based on the social interaction ratio, with a ratio less than 1 indicating susceptibility, and a ratio greater than 1 indicating resilience (Golden et al., 2011). In our previous work using the same strain of mice (McAllister et al., 2018), we found that many non-stressed controls had interaction ratios less than 1; therefore, we used a lower threshold of 0.5 for defining susceptibility. Using the same definition for the present analysis, we repeated the analysis of behaviour in the second social interaction test, excluding mice that were resilient to stress on the first test (4 WT-stress-vehicle, 4 KO-stress-vehicle, 4 KO-stress-fluoxetine). After doing so, there was a significant main effect of stress on time spent in the interaction zone [*F*(1,59) = 4.40, *p* = .040], with stress decreasing interaction time. However, excluding the resilient mice did not appreciably affect any other results (Supplemental Figure S1). There were no effects of fluoxetine that were specific to the stressed mice, and no differing effects of fluoxetine between genotypes.

Finally, to assess within-subjects changes in social interaction behaviour across the 4 weeks of fluoxetine treatment, we used the social interaction ratios to calculate difference scores for each mouse (Figure 5C), subtracting the observed ratio on the first social interaction test (t_1_) from the observed ratio on the second test (t_2_) (i.e., t_2_ - t_1_). A positive number thus represents an increase in the social interaction ratio. We again excluded mice that were found to be resilient to stress (interaction ratio • .5 at t_1_). Another mouse (KO-control-fluoxetine) was also excluded, due to a very small amount of time spent investigating the empty cage (3.9 s), resulting in an inflated interaction ratio. First, we assessed how interaction ratios changed across time in the non-stressed control groups, using one-sample t-tests to determine whether the observed mean difference score for each group differed from zero (a score of zero indicates no change over time). Surprisingly, in the vehicle-treated WT-control mice, interaction ratios increased significantly over time [*t*(10) = 2.92, *p* = .015]. With this exception, interaction ratios remained stable in the control groups [WT-control-fluoxetine: *t*(9) = 0.93, *p* = .376; KO-control-vehicle: *t*(8) = 0.29, *p* = .782; KO-control-fluoxetine: *t*(8) = 0.20, *p* = .845]. We next assessed how social interaction behaviour changed over time in the stressed mice. Consistent with our first hypothesis, in stressed mice that received vehicle treatment, interaction ratios did not change significantly over time in either WT [*t*(5) = 1.05, *p* = .340] or ZnT3 KO mice [*t*(4) = 1.79, *p* = .148]. But, contrary to our second hypothesis, treating stressed WT mice with fluoxetine did not increase interaction [*t*(9) = 1.57, *p* = .151]. Fluoxetine treatment also had no significant effect on stressed ZnT3 KO mice [*t*(5) = 2.06, *p* = .095], though it tended to increase interaction ratios. To summarize, in stress-susceptible mice, the effect of stress on interaction time persisted 4 weeks to the second social interaction test. However, our hypothesis that fluoxetine treatment would decrease social avoidance in WT mice, but not in ZnT3 KO mice, was not confirmed.

#### 3.2.2. Novelty-suppressed feeding

Anxiety-like behaviour in the NSF test was first assessed on day 4 post-RSD (Figure 6A). Three mice (1 KO-stress-vehicle, 2 KO-stress-fluoxetine) were excluded from this analysis (and analysis of the results from the second NSF test) because they did not respond well to the overnight food deprivation. These mice were cold to the touch and moved very little, or not at all, when placed in the novel open field. The mice were immediately withdrawn from the NSF test, placed on a heating pad, and provided with food.

**Figure 6.**
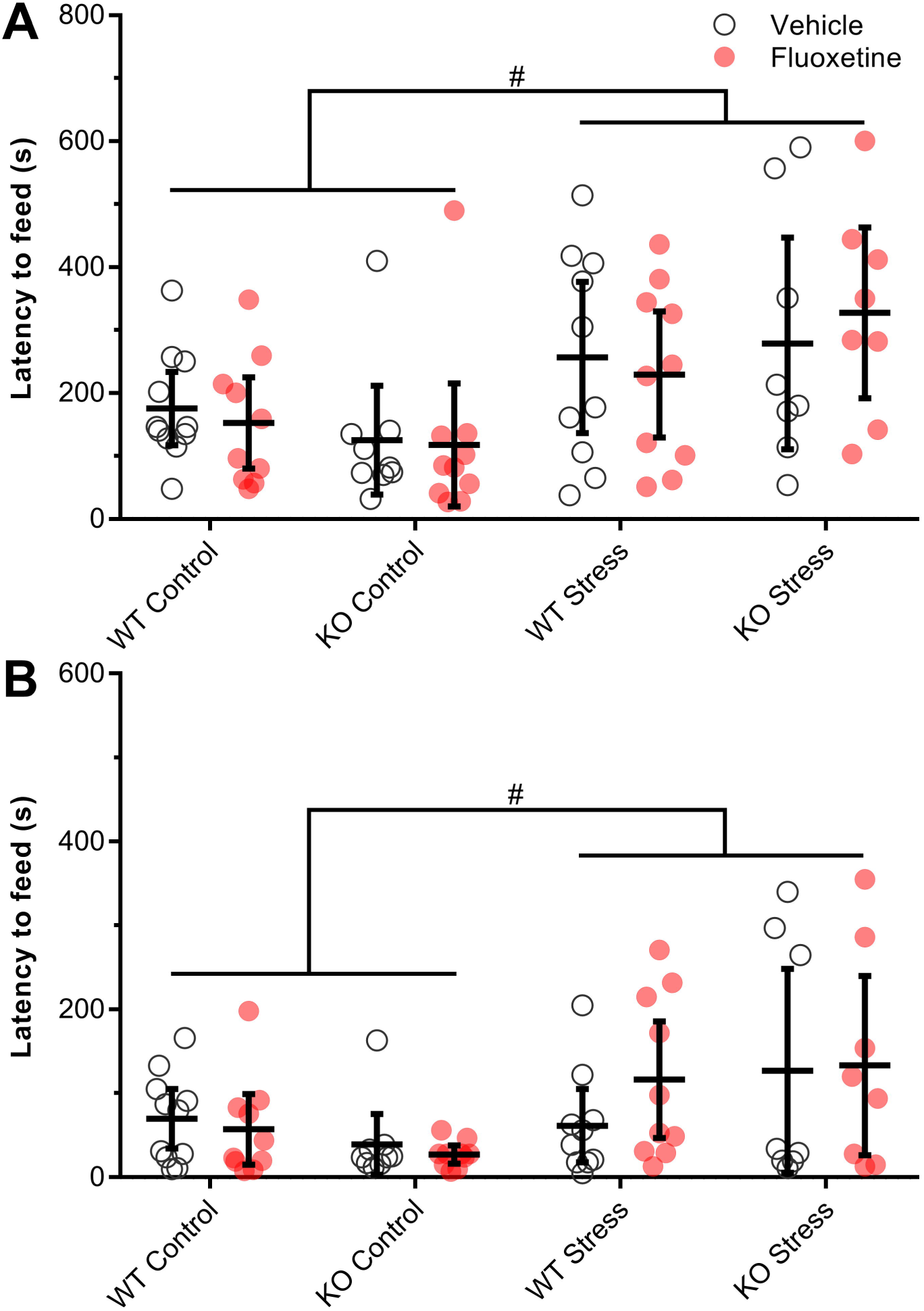
The effects of repeated social defeat stress, and subsequent fluoxetine treatment, on anxiety-like behaviour in the novelty-suppressed feeding (NSF) test. (**A**) Results from the first NSF test, after 3 days of fluoxetine treatment. Stress increased the latency to feed in a novel open field, indicating increased anxiety-like behaviour. As anticipated, fluoxetine had no effect after only 3 days of treatment. There was also no difference between genotypes. (**B**) Results from the second NSF test, after 24 days of fluoxetine treatment. The effect of stress on anxiety-like behaviour persisted to the second test. Contrary to our hypothesis, chronic fluoxetine treatment did not decrease anxiety-like behaviour. There was also no difference between genotypes. Error bars represent 95% CIs. ^#^effect of stress, *p* < .05

Stress increased the latency to feed in the novel field by 87% [*F*(1,68) = 16.25, *p* < .001], indicating increased anxiety-like behavior. There was no difference between genotypes [*F*(1,68) = 0.07, *p* = .787]. On the first NSF test, after only 3 days of fluoxetine treatment, we predicted that the drug would have no effect, which was confirmed [main effect of fluoxetine: *F*(1,68) = 0.01, *p* = .943]. There was no significant effect of stress on latency to feed in the home cage [*F*(1,68) = 3.12, *p* = .082], though the stressed mice did tend to take longer to feed (Table 2). The effect in the novel field, but not in the home cage, suggests that the increased latency to feed was caused by the anxiogenic environment of the novel field, rather than by more general effects of stress on feeding behaviour. In further support of this, there was no effect of stress on the amount of food consumed during the 5 min test in the home cage [*F*(1,68) = 1.16, *p* = .286; Table 2]. Interestingly, there was a significant interaction between fluoxetine treatment and genotype on food consumption [*F*(1,68) = 4.63, *p* = .035], with fluoxetine tending to increase consumption in the WT mice and decrease consumption in the ZnT3 KO mice. However, follow-up simple effects tests did not show a significant effect of fluoxetine treatment on either the WT [*F*(1,68) = 5.09, *p* = .054; Bonferroni-corrected] or ZnT3 KO mice [*F*(1,68) = 0.72, *p* = .400]. In summary, stress increased anxiety-like behaviour in the NSF test, regardless of genotype, and there was no effect of short-term fluoxetine treatment.

**Table 2.** Additional behavioural measures from the novelty-suppressed feeding tests. Statistics are reported as mean ± standard deviation. ^#^main effect of stress, *p* < .05 ^‡^main effect of drug, *p* < .05 *genotype × drug interaction, *p* < .05

|  | Vehicle |  |  |  | Fluoxetine |  |  |  |
| --- | --- | --- | --- | --- | --- | --- | --- | --- |
|  | Wild type |  | ZnT3 KO |  | Wild type |  | ZnT3 KO |  |
|  | Control | Stress | Control | Stress | Control | Stress | Control | Stress |
| **First test** | (n = 11) | (n = 10) | (n = 9) | (n = 8) | (n = 10) | (n = 10) | (n = 10) | (n = 8) |
| Latency to feed (s) – home cage | 31.1 ± 22.4 | 47.2 ± 31.1 | 33.9 ± 29.2 | 51.9 ± 57.3 | 37.3 ± 36.4 | 42.7 ± 33.0 | 26.6 ± 22.1 | 41.1 ± 26.7 |
| Weight loss (g)<sup>#</sup> | 2.7 ± 0.6 | 3.7 ± 0.8 | 2.9 ± 0.5 | 3.7 ± 0.9 | 2.8 ± 1.0 | 4.1 ± 1.2 | 3.3 ± 1.3 | 3.6 ± 0.4 |
| Consumption (g)<sup>*</sup> | 0.14 ± 0.06 | 0.14 ± 0.05 | 0.17 ± 0.06 | 0.15 ± 0.05 | 0.18 ± 0.05 | 0.17 ± 0.05 | 0.16 ± 0.07 | 0.13 ± 0.04 |
| **Second test** | (n = 11) | (n = 10) | (n = 9) | (n = 8) | (n = 10) | (n = 10) | (n = 10) | (n = 8) |
| Latency to feed (s) – home cage | 23.5 ± 21.6 | 23.6 ± 16.2 | 10.3 ± 6.8 | 41.5 ± 47.6 | 26.3 ± 20.5 | 26.2 ± 24.6 | 15.4 ± 8.6 | 16.5 ± 9.9 |
| Latency to feed (s) – diff. score<sup>#</sup> | -105.8 ± 73.7 | -195.2 ± 132.7 | -86.0 ± 66.8 | -151.9 ± 78.1 | -95.2 ± 113.8 | -113.1 ± 91.0 | -90.8 ± 127.5 | -194.0 ± 140.8 |
| Weight loss (g)<sup>#†</sup> | 2.8 ± 0.3 | 3.6 ± 0.4 | 2.8 ± 0.3 | 3.8 ± 0.4 | 2.2 ± 0.6 | 3.5 ± 0.5 | 2.4 ± 0.6 | 3.1 ± 0.7 |
| Consumption (g)<sup>†</sup> | 0.12 ± 0.04 | 0.16 ± 0.04 | 0.13 ± 0.07 | 0.11 ± 0.04 | 0.22 ± 0.05 | 0.17 ± 0.04 | 0.22 ± 0.07 | 0.22 ± 0.06 |

Anxiety-like behaviour in the NSF test was assessed a second time, 3 weeks after the first test. Difference scores, subtracting the latency to feed on the first test (t_1_) from the latency to feed on the second test (t_2_), showed that all groups had shorter latencies at t_2_, and that the decrease in latency was greater in the stressed mice [*F*(1,68) = 7.94, *p* = .006; Table 2], likely because they started from a much higher baseline at t_1_. Despite this, there was still a significant effect of stress on latency to feed at t_2_ [*F*(1,68) = 10.41, *p* = .002; Figure 6B], with the stressed mice exhibiting an 118% increase relative to non-stressed controls. There was also a trend toward an interaction between the effect of stress and the genotype of the mice [*F*(1,68) = 3.54, *p* = .064]. Follow-up simple effects tests showed that there was no effect of stress on the WT mice [*F*(1,39) = 1.29, *p* = .264; Welch’s test] but that stress increased latency to feed in the ZnT3 KO mice [*F*(1,33) = 8.18, *p* = .022; Welch’s test; Bonferroni-corrected]. This suggests that the effect of stress on anxiety-like behaviour may be more persistent in ZnT3 KO mice than in WT mice. However, because the interaction was not significant, this interpretation would require further experimental validation. As in the first NSF test, there was no significant effect of stress on the latency to feed in the home cage test [*F*(1,68) = 2.46, *p* = .121; Table 2]. Together, these results indicate that the effect of stress on anxiety-like behaviour persisted for at least 25 days after the final episode of RSD in mice of both genotypes.

On the second test, after more than 3 weeks of drug treatment, we anticipated that the stressed, fluoxetine-treated mice would exhibit less anxiety-like behaviour than the stressed, vehicle-treated mice, and that this effect might be specific to the WT mice. However, we did not observe a significant effect of fluoxetine treatment on latency to feed in the novel field [*F*(1,68) = 0.23, *p* = .632], and the effect of fluoxetine treatment did not differ based on genotype [stress × genotype interaction: *F*(1,68) = 0.41, *p =* .527]. Thus, our results provide no evidence that fluoxetine treatment reverses the anxiety-like behaviour induced by RSD stress.

Although fluoxetine treatment did not affect anxiety-like behaviour, 24 days of fluoxetine treatment did, unexpectedly, increase the amount of food consumed in the home cage test [*F*(1,68) = 39.93, *p* < .001; Table 2]. There was also a trend toward a three-way interaction between stress, genotype, and fluoxetine treatment on food consumption, though this was not significant [*F*(1,68) = 3.69, *p* = .059].

Finally, we examined differences in the amount of weight lost over the 16 h food restriction period prior to both the first and second NSF tests (Table 2). Previously, we found that RSD stress increased the amount of weight lost, possibly suggesting stress-induced metabolic changes (McAllister et al., 2018). Here, we also observed the same effect on the first NSF test [main effect of stress: *F*(1,68) = 15.75, *p* < .001], and this effect persisted at least 3 weeks to the second NSF test [*F*(1,68) = 69.00, *p* < .001]. These effects were also significant when weight loss was calculated as a percentage of body weight (data not shown). Interestingly, 3 days of fluoxetine treatment had no effect on weight loss in the first test [*F*(1,68) = 0.83, *p* < .365], but on the second test, after 24 days of treatment, there was a significant effect [*F*(1,68) = 17.99, *p* < .001], with fluoxetine decreasing the amount of weight lost over the food restriction period. Again, this effect was also significant when weight loss was calculated as a percentage of body weight (data not shown). In summary, while fluoxetine treatment did not affect anxiety-like behaviour in the NSF test, it may have had an effect on metabolism, reducing the amount of weight lost as a result of food restriction.

## 4. DISCUSSION

One goal of the present experiment was to assess whether neurogenesis can be modulated by stress in ZnT3 KO mice. Stress is commonly thought of as an inhibitor of hippocampal neurogenesis; in part, this fact has underlain the neurogenesis theory of depression since its inception (Jacobs, van Praag, & Gage, 2000). Focusing specifically on chronic social stress in mice, most previous reports indicate that stress inhibits neurogenesis, decreasing cell proliferation, survival, or neuronal differentiation (Chen, Huang, Hsu, 2015; Ferragud et al., 2010; Mitra et al., 2006; McKim et al., 2016; Schloesser, Lehmann, Martinowich, Manji, & Herkenham, 2010; Walker et al., 2015). It is interesting, then, that we observed a strong positive effect of stress on neurogenesis. Specifically, we labeled dividing cells by administering BrdU to mice 1 day after the final episode of RSD stress. When brains were collected 4 weeks later, and the number of BrdU^+^ cells in the SGZ and granule cell layer of the hippocampal dentate gyrus was counted, a substantial stress-induced increase was observed. Though unexpected, the observation of increased cell survival in response to stress is not unprecedented (De Miguel, Haditsch, Palmer, Azpiroz, & Sapolsky, 2018; Lagace et al., 2010). In particular, the present results are in alignment with those of Lagace et al. (2010), who – using a very similar, though more severe, RSD protocol – demonstrated an increase in the 4-week survival of cells born 1-day post-RSD. They also found that the increase in cell survival was limited to mice that were susceptible to social avoidance following stress, and that ablating neurogenesis by cranial x-ray irradiation made mice more resilient. We did not replicate the finding that a greater number of surviving cells is associated with susceptibility; in our experiment both susceptible and resilient mice had more BrdU^+^ cells than did the non-stressed controls, though we note that our definition of susceptible (interaction ratio < 0.5) differed from the definition used by Lagace et al. (2010) (interaction ratio < 1). The finding that blocking neurogenesis promotes resilience is also at odds with a recent report that the activity of adult-born granule cells, specifically in the ventral dentate gyrus, promotes resilience to social avoidance following RSD stress (Anacker et al., 2018).

Previously, we found that cell proliferation, as measured by immunolabeling for the endogenous proliferation marker Ki67, was not increased in the dentate gyrus of mice 1 day after the final episode of RSD stress (McAllister et al., 2018). It is important to note that these mice were subjected to a social interaction test prior to their brains being collected, just as the mice in the present study were subjected to the same test prior to the first BrdU injection. Thus, our previous finding provides evidence that the increased number of BrdU^+^ cells observed in the present study was due to an increase in the rate of cell survival, rather than an increase in proliferation at the time of the BrdU injections. It should be noted that, based solely on our own results, we cannot rule out the possibility that RSD stress increased blood-brain barrier (BBB) permeability, resulting in a greater concentration of BrdU reaching the hippocampus. This, rather than increased survival, could explain the increase in the number of BrdU^+^ cells observed 4 weeks later. However, Lagace et al. (2010) injected mice with BrdU 1-day post-RSD and killed them 2 h later to examine proliferation, and they found no effect of RSD stress on the number of BrdU^+^ cells (or Ki67^+^ cells) at this timepoint. This indicates that increased BrdU labeling due to increased BBB permeability is likely not a factor in these results, and it also supports our observation that proliferation is not increased 1-day post-RSD.

A limitation of the present experiment is that neuronal differentiation of BrdU^+^ cells was not examined. Adult-born cells in the dentate gyrus mostly develop into neurons (Encinas et al., 2006), but some instead become glial cells or do not express standard neuronal or glial markers. We did not quantify the percentage of BrdU^+^ cells that expressed a neuronal marker, so we do not know the proportion of surviving cells that became neurons. However, Lagace et al. (2010) found that the percentage of surviving BrdU^+^ cells that expressed the neuronal marker NeuN was consistent between stressed and control mice at approximately 75%. Others have shown that neuronal differentiation is not affected by chronic fluoxetine treatment (Encinas et al., 2006; Malberg et al., 2000; Santarelli et al., 2003). Together, these findings suggest that neuronal differentiation is unlikely to have been affected by stress or fluoxetine treatment in the present experiment.

In unpublished experiments from our laboratory, we previously observed that vesicular zinc is required for the pro-neurogenic effect, both on cell proliferation and survival, of chronic fluoxetine treatment in female mice (Boon, 2016). Based on this finding, a second goal of the present experiment was to test whether neurogenesis is affected by chronic fluoxetine treatment in male ZnT3 KO mice. The results, though not entirely conclusive, suggest it is not. We observed a positive effect of chronic fluoxetine treatment on the number of surviving cells, consistent with previous reports (Encinas et al., 2006; Santarelli et al., 2003). When we broke this comparison down by genotype, we found that fluoxetine significantly increased the number of surviving cells in WT mice but not in ZnT3 KO mice, as predicted. However, there was no significant interaction between the effect of fluoxetine treatment and the genotype of the mice; that is, fluoxetine treatment did not have a significantly greater effect on WT mice than it did on ZnT3 KO mice. When we assessed proliferation after 4 weeks of chronic fluoxetine treatment, we found that there was no effect in either genotype. There was a trend toward an effect of fluoxetine treatment in the non-stressed WT mice, but not in any of the other groups. This is not necessarily inconsistent with our hypothesis and with previous results; we anticipated that fluoxetine would not have an effect on neurogenesis in the ZnT3 KO groups, and social stress has previously been shown to have a suppressive effect on proliferation (Ferragud et al., 2010; Mitra et al., 2006; Walker et al., 2015), which may have counteracted any positive effect of fluoxetine in the stressed WT group. Thus, it would not be entirely surprising if only the non-stressed WT mice exhibited an effect of fluoxetine on proliferation. On the other hand, we observed no significant suppression of proliferation by stress; this does not necessarily preclude the possibility that stress counteracted the effects of fluoxetine, but it does make it seem less likely. Other researchers have similarly reported that social stress does not affect proliferation. Specifically, Schloesser et al. (2010) and McKim et al. (2016) both found that proliferation was unaffected by RSD stress, though these groups did find that stress inhibited other components of hippocampal neurogenesis (cell survival and neuronal differentiation, respectively). In summary, while our results are partially consistent with our previous findings that vesicular zinc is required for the pro-neurogenic effects of fluoxetine treatment, further validation – ideally in a larger sample providing more statistical power to detect an interaction effect, and possibly using a higher dosage of fluoxetine to produce a more robust effect on cell proliferation – will be required to more conclusively determine whether the results we previously observed in female mice also extend to males.

On the topic of cell proliferation, it is noteworthy that the effect of fluoxetine treatment on the non-stressed WT mice was quite variable in the present study, which may have contributed to a non-significant result in this group. This variability may be related to a limitation of the method used to administer fluoxetine. Fluoxetine was administered in the drinking water; in cages with more than one mouse (i.e., in the non-stressed groups), the dosage was calculated based on the average weight of the mice, and under the assumption that all mice drank an equal amount. Thus, if mice in a cage were of unequal weights, or if they consumed unequal volumes of water, then dosages of fluoxetine would have been somewhat variable between mice. The tradeoff for reduced accuracy of dosing is that administration through the drinking water eliminates any potential confounding effect of stress resulting from repeated daily injections.

A third goal of the present study was to serve as a replication of our previous finding that RSD stress causes social avoidance of a same-strain conspecific in WT mice, but not in ZnT3 KO mice, suggesting decreased susceptibility to stress in mice that lack vesicular zinc (McAllister et al., 2018). When we examined the effect of stress independently within each genotype, we observed that RSD stress significantly increased social avoidance (i.e., decreased interaction time and increased corner time) in WT mice but not in ZnT3 KO mice, consistent with our previous results. However, unlike our previous results, there was no significant interaction between the effect of RSD stress and the genotype of the mice on interaction time or corner time; that is, the effect of RSD stress on social avoidance was not significantly greater in the WT mice than in the ZnT3 KO mice. Thus, the present results provide some support for, but do not fully replicate, our previous findings, despite using samples of a similar size as in our previous experiment. It is possible that the size of the interaction effects may have been overestimated in our previous study, and that larger samples would be required to reliably detect these effect. Further replication will be required to test whether this is the case.

Finally, we assessed the effects of RSD stress, and subsequent treatment with fluoxetine, on the behaviour of WT and ZnT3 KO mice. Consistent with our previous findings (McAllister et al., 2018), RSD stress led to avoidance of an aggressive CD-1 mouse in the social interaction test and increased latency to feed in the NSF test, which can be interpreted as depression-like (i.e., social withdrawal) and anxiety-like behaviours, respectively. Furthermore, the anxiety-like behaviour persisted for at least 3 weeks, and, among mice that showed social avoidance (i.e., were defined as being susceptible to stress) on the first interaction test, social avoidance behaviour persisted for at least 4 weeks. The persistent nature of these effects allowed us to examine whether they could be reversed by chronic treatment with fluoxetine. Contrary to several previous findings (David et al., 2009; Santarelli et al., 2003; Surget et al., 2008; Wang et al., 2008), we could not detect an effect of fluoxetine on anxiety-like behaviour in the NSF test. In the social interaction test with an aggressive CD-1 mouse, fluoxetine had a moderate effect, decreasing the amount of time spent in the corners (i.e., decreasing social avoidance), but not significantly increasing the time spent interacting (i.e., in close proximity) with the CD-1 target. This is broadly consistent with findings that chronic fluoxetine treatment decreases social avoidance following RSD stress in mice (Berton et al., 2006; Tsankova et al., 2006; Vialou et al., 2015; but see Venzala, García-García, Elizalde, Delagrande, & Tordera, 2012), though the effect may differ across strains (Razzoli et al., 2011). It should be noted, though, that the effect of fluoxetine in the present experiment was a general decrease in social avoidance across all groups, rather than an antidepressant-like effect that was specific to the stressed mice. Overall, the behavioural results provided no support for our hypothesis that ZnT3 KO mice would fail to benefit from the anti-anxiety and antidepressant-like effects of fluoxetine – though the lack of robust effects of fluoxetine treatment, even in WT mice, make it difficult to draw any strong conclusions regarding this hypothesis. In the future, this hypothesis might be better tested by using a higher dosage of fluoxetine, or perhaps a different antidepressant drug that exerts stronger behavioural effects.

Interestingly, the most robust effect of fluoxetine treatment in the present experiment was to attenuate the amount of weight lost over the 16 h food deprivation period prior to the NSF test. This was the opposite of the effect of RSD stress, suggesting that chronic fluoxetine treatment may help to decrease the metabolic abnormality caused by RSD – though the exact nature of this abnormality remains to be uncovered. Notably, increased caloric intake following RSD stress has been linked to increased ghrelin signaling (Lutter et al., 2008), and fluoxetine treatment has been found to reverse the stress-induced increase in food intake, while also altering the circadian profile of ghrelin levels (Kumar et al., 2013).

The broad objective of the present experiment was to address two main questions. The first was whether mice that lack vesicular zinc would exhibit modulation of adult hippocampal neurogenesis by RSD stress. The second was whether hippocampal neurogenesis in these mice could be modulated by chronic fluoxetine treatment, and whether neurogenesis would be required for any anti-anxiety or antidepressant-like effects of the drug. The answer to the first question was clear. In both WT and ZnT3 KO mice, RSD stress clearly increased the number of surviving cells born 1 day after the final episode of defeat, clearly indicating that at least this aspect of neurogenesis can be modulated by stress in ZnT3 KO mice. Regarding the second question, we found no evidence that the behavioural effects of fluoxetine were absent, or in any way altered, in mice that lack vesicular zinc – though the lack of strong behavioural effects of fluoxetine even in WT mice did not provide an ideal test of this hypothesis. Finally, the results provided partial support for our previous findings that vesicular zinc is required for the neurogenic effect of chronic fluoxetine treatment, and that a lack of vesicular zinc modifies the behavioural effects of social defeat stress.

## Acknowledgements

The authors thank Nicoline Bihelek for her assistance with the behavioural testing. This research was supported by an NSERC Discovery Grant (RHD) and by doctoral scholarships from NSERC and the Killam Trusts (BBM). The data that support the findings of this study are available from the corresponding author upon reasonable request.

