## Supplemental methods, data and tables for "Effects of social defeat stress and fluoxetine treatment on neurogenesis and behaviour in mice that lack zinc transporter 3 (ZnT3) and vesicular zinc"

Immunofluorescence Labeling

The procedure for BrdU immunofluorescence was as follows: 1 h in 2 M hydrochloric acid with gentle agitation every 15 min; 6 × 15 min wash in PBS; overnight incubation at room temperature in PBS containing 0.3% Triton X-100 (PBSx) with 2% normal goat serum (NGS) and the primary antibody (1:200 rat anti-BrdU, Bio-Rad MCA2060); 3 × 10 min wash in PBS; overnight incubation at room temperature in PBS containing the secondary antibody (1:500, biotin-conjugated goat anti-rat, Jackson ImmunoResearch 112-065-167); 3 × 10 min wash in PBS; 1 h incubation in PBS containing the tertiary antibody (1:500, Alexa-Fluor 594-conjugated streptavidin, Jackson ImmunoResearch 016-580-084) with 0.1 µg/ml of 4′,6-diamidino-2-phenylindole (DAPI; Sigma) added for the final 15 min; 3 × 10 min wash in PBS.

For Ki67 immunofluorescence labeling, the procedure was as follows: 3 × 10 min wash in PBS; blocking for 1 h in PBSx with 4% NGS; incubation overnight at room temperature in PBSx containing 2% NGS and the primary antibody (1:2000, 0.5 μg/ml, rabbit anti-Ki67, Abcam, ab15580); 3 × 10 min wash in PBS; incubation overnight at room temperature in PBSx containing the secondary antibody (1:1000, biotin-conjugated goat anti-rabbit, Jackson ImmunoResearch 111-065-144); 3 × 10 min wash in PBS; incubation for 1 h in PBS containing the tertiary antibody (1:500, Alexa-Fluor 594-conjugated streptavidin, Jackson ImmunoResearch 016-580-084) with 0.1 µg/ml of DAPI added for the final 15 min; 3 × 10 min wash in PBS.

Following immunofluorescence labeling, sections were mounted to gelatin-coated slides, coverslipped with fluorescence mounting medium, and stored in the dark at 4 °C until analysis.

SUPPLEMENTAL FIGURES

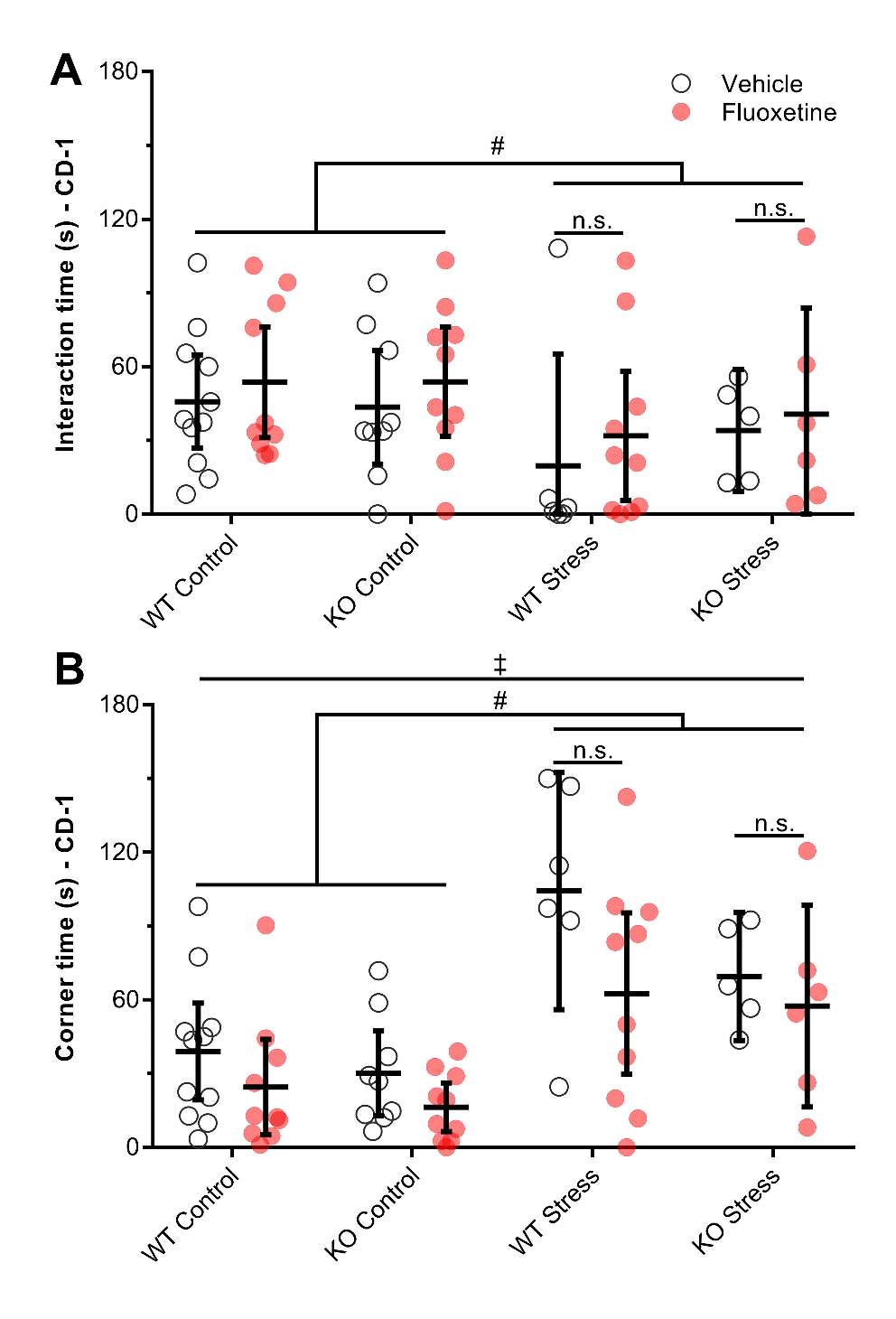

**Figure S1.** Results from the second social interaction test, excluding susceptible mice (interaction ratio ≥ 0.5 in the first social interaction test). (A) Stress decreased interaction time with a novel, aggressive CD-1 mouse. Fluoxetine did not have a counteracting effect, either overall or specifically in stressed WT or ZnT3 KO mice. (B) Stress increased corner time in the presence of a CD-1 mouse; together with decreased interaction time, this indicates that social avoidance persisted at least 4 weeks. Fluoxetine decreased corner time, indicating decreased social avoidance, though this effect was not limited to the stressed mice. Error bars represent 95% CIs. ^#^effect of stress, *p* < .05; ^‡^effect of fluoxetine, *p* < .05; n.s. indicates *p* > .05

SUPPLEMENTAL TABLES

| **Table S1.** ANOVA results (effects of genotype and stress) from the first social interaction test. The test was conducted prior to the start of fluoxetine treatment. | | | | |
| --- | --- | --- | --- | --- |
|  | D.F. | Genotype | Stress | Genotype × stress |
| **First social interaction test** |  |  |  |  |
| Interaction time - conspecific | 1, 75 | *F* = 1.00, *p* = .321 | ***F* = 5.23, *p* = .025** | *F* = 1.36, *p* = .248 |
| Corner time - conspecific | 1, 75 | *F* = 0.51, *p* = .476 | ***F* = 8.80, *p* = .004** | *F* = 0.33, *p* = .568 |
| Interaction time - CD-1 | 1, 75 | *F* = 1.42, *p* = .238 | ***F* = 34.70, *p* < .001** | *F* = 1.59, *p* = .212 |
| Corner time - CD-1 | 1, 75 | *F* = 0.10, *p* = .750 | ***F* = 30.18, *p* < .001** | *F* = 1.93, *p* = .169 |
| Interaction ratio – CD-1 | 1, 75 | *F* = 3.28, *p* = .074 | ***F* = 14.71, *p* < .001** | *F* = 0.52, *p* = .472 |
| Interaction time – empty cage | 1, 75 | *F* = 3.07, *p* = .084 | ***F* = 8.28, *p* = .005** | *F* = 0.36, *p* = .549 |
| Distance – empty cage | 1, 75 | *F* = 0.99, *p* = .323 | *F* = 1.56, *p* = .216 | *F* < 0.01, *p* = .997 |

| **Table S2.** ANOVA results (effects of genotype, stress, and fluoxetine). Results are from the second social interaction test, after 4 weeks of fluoxetine treatment; from the first and second novelty-suppressed feeding (NSF) tests, after 3 days and 24 days of fluoxetine treatment, respectively; and from analysis of adult hippocampal neurogenesis following fluoxetine treatment. | | | | |
| --- | --- | --- | --- | --- |
|  | D.F. | Genotype | Stress | Fluoxetine |
| **Social interaction test** |  |  |  |  |
| ***All mice*** |  |  |  |  |
| Interaction time – CD-1 | 1, 71 | *F* = 0.13*, p* = .722 | *F* = 2.80*, p* = .098 | *F* = 0.71*, p* = .403 |
| Corner time – CD-1 | 1, 71 | *F* = 1.51*, p* = .223 | ***F* = 29.27*, p* < .001** | ***F* = 5.59*, p* = .021** |
| ***Susceptible only*** |  |  |  |  |
| Interaction time – CD-1 | 1, 59 | *F* = 0.40*, p* = .529 | ***F* = 4.40*, p* = .040** | *F* = 1.22*, p* = .275 |
| Corner time – CD-1 | 1, 59 | *F* = 3.09*, p* = .084 | ***F* = 32.20*, p* < .001** | ***F* = 6.41*, p* = .014** |
| CD-1 Interaction Ratio – diff. score | 1, 58 | *F* = 0.74*, p* = .393 | ***F* = 4.12*, p* = .047** | *F* = 0.08*, p* = .779 |
| **NSF test** |  |  |  |  |
| ***First test*** |  |  |  |  |
| Latency to feed – novel field | 1, 68 | *F* = 0.07*, p* = .787 | ***F* = 16.25*, p* < .001** | *F* = 0.01*, p* = .943 |
| Latency to feed – home cage | 1, 68 | *F* = 0.03*, p* = .876 | *F* = 3.12*, p* = .082 | *F* = 0.29*, p* = .595 |
| Weight loss | 1, 68 | *F* = 0.12*, p* = .735 | ***F* = 15.75*, p* < .001** | *F* = 0.83*, p* = .365 |
| Food consumption | 1, 68 | *F* = 0.22*, p* = .643 | *F* = 1.16*, p* = .286 | *F* = 0.82*, p* = .369 |
| ***Second test*** |  |  |  |  |
| Latency to feed – novel field | 1, 68 | *F* = 0.07*, p* = .787 | ***F* = 16.25*, p* < .001** | *F* = 0.01*, p* = .943 |
| Latency to feed – home cage | 1, 68 | *F* = 0.59*, p* = .445 | *F* = 2.46*, p* = .121 | *F* = 0.50*, p* = .484 |
| Weight loss | 1, 68 | *F* = 0.07*, p* = .790 | ***F* = 69.00*, p* < .001** | ***F* = 17.99*, p* < .001** |
| Food consumption | 1, 68 | *F* = 0.24*, p* = .624 | *F* = 0.38*, p* = .542 | ***F* = 39.93*, p* < .001** |
| Latency to feed – difference score | 1, 68 | *F* = 0.02*, p* = .892 | ***F* = 7.94*, p* = .006** | *F* = 0.22*, p* = .642 |
| **Neurogenesis** |  |  |  |  |
| BrdU^+^ cell count (survival) | 1, 71 | *F* = 0.38*, p* = .540 | ***F* = 21.88*, p* < .001** | ***F* = 5.64*, p* = .020** |
| Ki67^+^ cell count (proliferation) | 1, 71 | *F* = 0.11*, p* = .747 | *F* = 0.09*, p* = .769 | *F* = 1.26*, p* = .265 |

| **Table S2 (continued).** | | | | |
| --- | --- | --- | --- | --- |
|  | Genotype × stress | Genotype × drug | Stress × drug | Genotype × stress × drug |
| **All mice** |  |  |  |  |
| Interaction time – CD-1 | *F* = 0.25*, p* = .622 | *F* = 0.27*, p* = .604 | *F =* 0.13*, p* = .716 | *F =* 0.13*, p* = .721 |
| Corner time – CD-1 | *F* = 0.01*, p* = .905 | *F* = 0.01*, p* = .912 | *F =* 0.28*, p* = .600 | *F <* 0.01*, p* = .951 |
| **Susceptible only** |  |  |  |  |
| Interaction time – CD-1 | *F* = 0.57*, p* = .452 | *F* = 0.01*, p* = .927 | *F <* 0.01*, p* = .986 | *F* = 0.06*, p* = .813 |
| Corner time – CD-1 | *F* = 0.49*, p* = .486 | *F* = 0.88*, p* = .351 | *F =* 0.62*, p* = .435 | *F* = 0.80*, p* = .376 |
| CD-1 Interaction Ratio – diff. score | ***F* = 7.18*, p* = .010** | ***F* = 6.53*, p* = .013** | ***F* = 6.35*, p* = .014** | *F* = 0.05*, p* = .826 |
| **NSF test** |  |  |  |  |
| ***First test*** |  |  |  |  |
| Latency to feed – novel field | *F* = 2.50*, p* = .118 | *F* = 0.50*, p* = .482 | *F* = 0.16*, p* = .692 | *F* = 0.22*, p* = .644 |
| Latency to feed – home cage | *F* = 0.13*, p* = .720 | *F* = 0.42*, p* = .520 | *F* = 0.22*, p* = .644 | *F* = 0.06*, p* = .813 |
| Weight loss | *F* = 2.43*, p* = .124 | *F* = 0.05*, p* = .831 | *F* = 0.01*, p* = .922 | *F* = 1.05*, p* = .310 |
| Food consumption | *F* = 0.45*, p* = .503 | ***F* = 4.63*, p* = .035** | *F* = 0.15*, p* = .699 | *F* = 0.07*, p* = .788 |
| ***Second test*** |  |  |  |  |
| Latency to feed – novel field | *F* = 3.54*, p* = .064 | *F* = 0.41*, p* = .527 | *F* = 1.27*, p* = .263 | *F* = 0.42*, p* = .522 |
| Latency to feed – home cage | *F* = 2.45*, p* = .122 | *F* = 1.52*, p* = .222 | *F* = 2.17*, p* = .146 | *F* = 2.10*, p* = .152 |
| Weight loss | *F* = 0.50*, p* = .482 | *F* = 0.42*, p* = .520 | *F* = 0.10*, p* = .751 | *F* = 2.71*, p* = .104 |
| Food consumption | *F* = 0.35*, p* = .555 | *F* = 2.50*, p* = .118 | *F* = 2.30*, p* = .134 | *F* = 3.69*, p* = .059 |
| Latency to feed – difference score | *F* = 0.40*, p* = .531 | *F* = 2.03*, p* = .159 | *F* = 0.12*, p* = .729 | *F* = 1.23*, p* = .271 |
| **Neurogenesis** |  |  |  |  |
| BrdU^+^ cell count (survival) | *F* = 0.84*, p* = .362 | *F* = 0.53*, p* = .470 | *F* = 0.21*, p* = .649 | *F* = 0.42*, p* = .520 |
| Ki67^+^ cell count (proliferation) | *F* = 0.09*, p* = .769 | *F* = 1.77*, p* = .188 | *F* = 0.91*, p* = .344 | *F* = 1.07*, p* = .304 |
